# Canonical babbling during vocal turn taking and independent vocal play

**DOI:** 10.1101/2020.10.09.333872

**Authors:** Helen L. Long, Gordon Ramsay, Dale D. Bowman, Megan M. Burkhardt-Reed, D. Kimbrough Oller

**Affiliations:** Waisman Center, University of Wisconsin-Madison, Madison, WI, USA; Marcus Autism Center, Children’s Healthcare of Atlanta, Atlanta, Georgia, USA; Department of Pediatrics, Emory University School of Medicine, Atlanta, Georgia, USA; Center for Translational Social Neuroscience, Emory University, Atlanta, Georgia, USA; Department of Mathematics, University of Memphis, Memphis, Tennessee, USA; Institute for Intelligent Systems, University of Memphis, Memphis, Tennessee, USA; School of Communication Sciences and Disorders, University of Memphis, Memphis, Tennessee, USA; Konrad Lorenz Institute for Evolution and Cognition Research, Klosterneuburg, Austria

**Keywords:** vocal development, social motivation, exploratory vocalization, endogenous activity, infant behavior, autism spectrum disorder

## Abstract

There is a growing body of research emphasizing the role of social and endogenous motivations in human development. The present study evaluated canonical babbling across the second-half year of life using all-day recordings of 98 infants participating in a longitudinal study. Canonical babbling ratios were calculated from human coding along with Likert-scale ratings on vocal turn taking and vocal play in each segment. Ratios across all infants were significantly elevated during high turn taking and also during high vocal play. We conclude that both social and endogenous motivations may drive infants’ tendencies to produce their most advanced vocal forms.

## Introduction

Canonical babbling has been long established as a robust stage of prelinguistic vocal development, occurring prior to the emergence of early words, having been argued to constitute a necessary foundation for vocabulary development [1–3]. To our knowledge, there is no published research evaluating the role of exploratory motivation in infants’ production of canonical babbling and no direct evaluation of the extent to which social engagement in vocal turn taking affects it. In the present research, we observed babbling in naturalistic settings recorded longitudinally for a sample of infants, half of whom had been identified as being at low risk and half at high risk for autism. Segments extracted from all-day home audio recordings were rated for levels of infant turn taking and independent vocal play to measure the degree of social and non-social vocal activity (and thus, social and exploratory motivations, respectively). We examined these findings within an evolutionary developmental biology (evo-devo) framework [4–7], in part to inform our understanding of how babbling may be used to signal wellness and developmental progress to caregivers [8–10].

### Canonical babbling and social interaction

Previous research examining the emergence of infant vocalizations deemed to be precursors to speech (hereafter, protophones), including canonical babbling, has largely focused on interactions with parents, rather than on internal motivations of the infant [11–20]. There is a growing body of research, however, examining intrinsically motivated vocal activity in infancy [21–23]. When evaluating the emergence of canonical babbling, there is reason to consider potential differences in the motivations underlying production of prelinguistic protophones across social and nonsocial contexts. We and others have hypothesized positive selection pressure on the production of infant protophones [21,24,25]. Infant protophones [2] are considered foundations for speech in that they manifest a capacity to produce phonation and other speech characteristics voluntarily, in the absence of external stimulation, as is required in speech. Such baby sounds can be seen as fitness signals, selected because they tend to elicit long-term investment from caregivers, required across the lengthy period of relative helplessness, or altriciality, of infant humans [8,21,26,27]. Parents throughout hominin history can be viewed as having implemented the selection of the infant trait of endogenous protophone production. In accord with the fitness signaling hypothesis, not only the quantity, but also the *quality* of infant vocalizations is considered a salient and reliable signal of fitness.

In line with this reasoning, it might be seen as advantageous for infants to produce their most advanced vocal forms during periods of maximal caregiver attention (i.e., during social interaction). Empirical evidence has been presented to show that caregivers are keenly aware of their infants’ developmental capabilities, including their vocal capabilities [28–30]. Higher rates of canonical syllables (as opposed to less advanced protophones) during social interaction than during periods of aloneness could suggest a social motivation for producing more advanced protophones. If the idea is on target, we might propose that canonical babbling was selected as a salient signal of developmental progress, especially during social interaction. Thus, infants with hypothetically reduced social motivation, such as those at high risk for autism [31–33] might be expected to produce lower rates of canonical babbling during social interaction than would occur in typically developing infants.

### On the role of exploratory vocal play in infant vocal development

Contradicting the seemingly implicit assumption in child development literature that infant vocalizations are routinely produced during social interaction, several researchers have recently emphasized the role of intrinsic motivation in the development of emotional and cognitive systems, including those related to vocal development [22,23,34]. We previously showed that typically developing infants produce protophones (both canonical and precanonical) predominantly endogenously, that is, with no indication of social directivity [21]. Even in laboratory recordings, during periods when parents are instructed to seek social interaction with infants, most protophones (∼60%) appear not to be directed to them. This predominance of endogenous vocalization is even stronger (∼80%) when parents are present with infants but not attempting to engage them. The numbers of protophones produced by human infants are prodigious, estimated based on randomly-sampled segments from all-day recordings, at 4-5 per minute every waking hour across the entire first year of life, and occurring at least 5 times more frequently than crying even from the first month of life [27,35]. Because protophone production is overwhelmingly produced in the absence of social interaction, and often when infants are alone in a room, it is hard to escape the impression that these vocalizations are internally motivated as a special kind of play that occurs very frequently in human infants, but rarely if ever in other ape infants [36].

Play is well established to be important throughout development [34,37–41]. Stark described vocal play as highly variable, with infants producing sounds in new and repeated combinations, modifying patterns and features during bouts of independent infant vocal activity [3]. During vocal play we see infants exploring sensorimotor aspects of the vocal apparatus and practice with various properties of sounds such as syllabic structure, amplitude, and pitch control. This description evokes the notion that vocal play can be considered a sensorimotor exploration of the vocal mechanism, which may be necessary to learn and master speech production. Thus, it is reasonable to expect infants to increase their production of canonical syllables during bouts of independent vocal play as they explore the sensorimotor aspects of sounds within their repertoires or perhaps even expand their repertoires as a function of exploration [42].

### Hypotheses

These results highlight a need for additional research on infant vocal tendencies across social and nonsocial circumstances. Specifically, we need to consider frequency of occurrence of advanced infant vocal forms (in particular, canonical babbling) during 1) vocal turn taking, which would suggest socially-motivated babbling, and 2) independent protophone production, which we shall refer to here as *vocal play* [3,43], and which would suggest endogenous motivation for vocal activity. Our assumption is that both social and exploratory motivations are involved in the production of protophones—even during social interaction, one can imagine that playful exploration of sounds and perhaps relations of the protophones with the sounds of the interacting other speakers may motivate infant activity. And it is especially important to determine the role of canonical babbling, because it is so much like mature speech, during these different circumstances.

The present research, thus, compares infant canonical babbling ratios (CBRs, number of canonical syllables divided by number of all protophone syllables) during recorded segments across periods with high and low frequencies of turn taking (TT) and periods with high and low frequencies of vocal play (VP). The research focuses on three ages during the second half-year of life—earlier protophones are almost entirely non-canonical, so we avoid the floor effect of CBRs before 6 months. We first evaluated possible differences in CBR between infants of high and low maternal education level, treated as a proxy for socioeconomic status (SES), and found no significant differences; therefore, we do not report maternal education effects in the data below. Sex differences were evaluated in a recent study from our laboratory using the present dataset, and no significant sex differences for CBR were found [44]; therefore, we also do not include sex as a variable in the present work.

*Age:* Based on extensive prior research on this topic [3,24,45–47], we predicted:

1. Higher CBRs will occur at the later ages than earlier ages, highlighting infants’ increasing ability to control the speech mechanism as they develop. *Risk:* Because our sample was selected from a larger, longitudinal study on vocal development in infants with and without autism risk, we had the opportunity to examine CBR differences across the groups to evaluate whether hypothetically different levels of social motivation significantly impact the overall production of canonical syllables. We predicted:
2. Higher CBRs will occur in the LR group compared to the HR group, a prediction based on the predominant, albeit inconsistent findings of the existing literature on canonical babbling in infants at risk for or diagnosed with autism [48]. *Turn Taking and Vocal Play:* Our primary analyses of interest addressed Turn Taking (TT) and Vocal Play (VP) separately. Specifically, we predicted:
3. Higher CBRs will occur during segments with high TT compared to low TT, a prediction based on the existing literature showing positive effects of social interaction on infant vocal development [12,15,49], and
4. Higher CBRs will occur during segments with high VP compared to low VP, a prediction based on the growing body of evidence indicating high rates of endogenous vocal activity proposed to signal fitness to caregivers [21,24].

## Materials and Methods

The institutional review boards of the University of Memphis and Emory University approved the procedures used in this study. Families provided written consent prior to participation.

### Participants

As part of an NIH-funded Autism Center of Excellence conducted at the Marcus Autism Center in Atlanta, Georgia, 100 families of newborn infants were recruited via flyers, advertisements, social media and community referrals to participate in a longitudinal sibling study of development across the first three years of life. We analyzed data from 98 infants (two infants did not complete recordings at the ages studied). Infants were recruited as being either at high risk (HR, n=49) or low risk (LR, n=49) for autism. Infants were deemed HR if they had at least one older biological sibling with a confirmed autism diagnosis, and LR if they had no familial history of autism in 1^st^, 2^nd^, or 3^rd^ degree relatives. Sex and self-reported maternal education (as a proxy for SES) measures were balanced to the extent possible in accord with known autism male-to-female ratios [50] and SES make-up of participants living in the greater Atlanta area who were willing and able to participate in a 3-year longitudinal study. Low/High SES groups were based on a median split of maternal education in the entire cohort. Table 1 presents demographic information for the infants included in this study.

**Table 1.** Participant demographics.

legend: Numbers of infants by Risk, Sex, and maternal education as a proxy for socioeconomic status (SES). \*One infant's family did not report maternal education.
|  |  | **High-Risk** | **Low-Risk** | **Total** |
| --- | --- | --- | --- | --- |
| **Total** |  | **49** | **49** | **98** |
| **Sex** | **Male** | 34 | 30 | 64 |
|  | **Female** | 15 | 19 | 34 |
| **SES\*** | **Low SES** | 26 | 18 | 44 |
|  | **High SES** | 22 | 31 | 53 |

Families were asked to complete audio recordings once a month from 1-36 months of age. This study used data collected between 6.5 and 13 months of age to represent the typical range of expected onset for and high infant activity in canonical babbling. These data were grouped into three age ranges for analysis and labeled with reference to the approximate mean age within each group: 6.5-8.49 months (7.5 months), 8.5-10.49 months (9.5 months), and 10.5-13 months (12 months). It should be noted that the 12-month age group included a slightly larger age range (2.5 months) than the 7.5- and 9.5-months age groups (2 months). There were no significant within-age-group differences between risk groups, as shown in Table 2.

**Table 2.** Participant age groups.

legend: Descriptive statistics (mean age, age range) for the three age groups, 7.5, 9.5, and 12 months, separated by risk group, LR and HR.
| **Age Group** | **Risk Group** | **Mean Age** | **Age Range (SD)** |
| --- | --- | --- | --- |
| **7.5** | **LR** | 7.50 | 6.51 – 8.48 (.59) |
|  | **HR** | 7.53 | 6.51 – 8.48 (.59) |
| **9.5** | **LR** | 9.41 | 8.52 – 10.42 (.54) |
|  | **HR** | 9.53 | 8.52 – 10.49 (.57) |
| **12** | **LR** | 11.64 | 10.52 – 12.92 (.69) |
|  | **HR** | 11.65 | 10.52 – 12.92 (.67) |

### Audio Recordings

Audio recordings were completed using LENA recording devices [51,52]. These devices are battery powered and secured inside the pocket of a special vest or clothing item with button clasps and can record up to 16 hours of audio per charge. LENA devices have a 16 kHz sampling rate and given the low mouth-to-microphone distance, usually offer excellent audio quality for human coding of recorded material.

#### Recording procedures

Families completed all-day recordings starting from the first month. Once a month, parents were provided with a LENA recording device and were supplied regularly with appropriately sized clothing for their infant to wear throughout the day, as well as full instructions on how to carry out recordings. The device was returned to the research project staff at the Marcus Autism Center each month following recording days for data processing. Each family completed ∼5 total recordings (range: 1-7) across the ages studied, with an average recording time of approximately 11 hours per day.

### Coding procedures

Twenty-one 5-minute segments were randomly extracted from each recording and coded in real-time for infant utterance counts by 16 trained graduate student coders in the Origin of Language Laboratories (OLL) at the University of Memphis. Graduate student coders were trained to differentiate canonical and non-canonical syllables during real-time coding and to rate the extent to which infants produced socially interactive (TT) and endogenous (VP) vocalizations during completion of the questionnaire that was filled out at the end of coding of each 5-minute segment. The six-week training procedure is detailed in Oller et al. [27]. OLL staff were blinded to all diagnostic and demographic information associated with each infant recording throughout the coding process. From these 21, eight segments, all having been coded as showing high volubility (infant protophone rate) and a range from high to low infant-directed speech were selected from each recording, yielding a total of 3799 segments. Fifteen of these segments were later excluded on the basis of having no infant vocalizations (in all of the 21 segments for that day, the infant produced no protophones); therefore, final analyses were completed on a total of 3,784 segments. The amount of infant-directed speech (IDS) was rated using the questionnaire that followed each of the 21 segments coded in the first coding pass. The questionnaire was also used to indicate environmental contextual factors for each segment, including audibility, other-person activity level, and aloneness of the infant. For both coding passes, each questionnaire item required a 5-point Likert-scale response to the relevant question, e.g., for IDS, “How often did someone talk to the infant?”

#### Canonical babbling ratios

In a second pass of coding, the 8 selected segments were coded in real-time for infant canonical and non-canonical syllables. Listeners identified a total of 30,263 canonical syllables, and 233,877 noncanonical syllables across the segments. To measure the emergence of advanced vocal forms, a *canonical babbling ratio* (CBR) was calculated as the total number of canonical syllables divided by the total number of syllables in each segment. Means and standard deviations of CBRs were calculated for each infant at each age. Occasionally, families did not complete a monthly recording, and for those cases there were no data at the infant’s age to include in the analysis. In cases where there were multiple recordings within an age for an infant, the means and SDs of these recordings were averaged prior to analysis.

#### Turn Taking and Vocal Play

Following syllable coding of each 5-minute segment, coders answered a 17-item questionnaire regarding how often infants used vocalizations for various functions based on the audible context of the infant’s environment in each segment. See Long et al. [21] for theoretical perspectives on making intuitive judgments of infant vocal functions. We used two items from the questionnaire to broadly assess rate of vocalizations judged to be social or exploratory within each segment. Specifically, the relevant questions were:

1. Were any of the infant’s protophones used in vocal turn taking with another speaker?
2. Were any of the infant’s protophones purely vocal play or vocal exploration?

Coders were trained on the identification of a variety of infant vocal functions including the two items listed. *Turn-Taking* was defined to refer to infant protophones produced during audible back and forth of vocalizations (and/or speech in the case of a verbal partner) between the infant and another person (i.e., caregiver, sibling, etc.); turn-taking in this definition requires that the infant be perceived as responding to infant-directed speech, not merely vocalizing during caregiver talk, which is often directed to other speakers.

*Vocal Play* was defined, as inspired by Stark [3], to refer to infant protophones produced independent of social interaction or any attempt on the part of the infant to initiate an interaction, as in calling to the caregiver. Often in vocal play, the infant appears to be manipulating vocalic (pitch level, pitch change, and loudness) and even consonant-like elements without any social intention. The vocalizations of vocal play must be judged to be of sufficient magnitude to be noticed and to have been voluntarily produced (not accidentally, as in the case of effort grunts).

After listening to each segment, coders responded to each question using a Likert Scale which aligned to the following rating designations: *1 = Never, 2 = Less than half the time, 3 = About half the time, 4 = More than half the time, 5 = Close to the whole time*. For example, a TT rating of 5 was applied to segments where a caregiver was clearly speaking to the infant, and the infant was protophones in an apparent back and forth vocal interaction for essentially the whole segment. Segments with a VP rating of 5 would indicate the listener perceived the vast majority of infant protophones across the segment as playful and exploratory and not directed to another person in any way. The subjective Likert Scale response system is intended to simulate the kinds of judgments caregivers make (we presume often unconsciously) regarding the function of vocalizations produced by infants throughout the day.

Vocal play and turn-taking, in accord with our definitions, are in principle independent of each other—no infant utterance can properly be treated as both, according to the definitions. Thus, vocal play in our definition is not socially directed and excludes socially interactive playful vocalization. Thus, any protophone produced in vocal interaction was deemed TT, but not VP, even if the interaction was thought to be playful. In addition, some protophones produced in the absence of vocal interaction, especially those produced with low intensity, were judged as pertaining to neither TT nor VP.

Although TT and VP are in principle mutually exclusive, a particular segment could include both TT and VP at different points during the segment. We have previously discussed the potential for vocal play to be a catalyst for parent-initiated social interaction [21]. If, for example, the infant is playing with the characteristics of a new sound, parents may repeat these sounds to the infant, initiating a back-and-forth vocal sequence. Thus, in segments with high infant vocal activity containing both interactive and non-interactive protophones, a segment could be coded as having high TT and high VP. This happened rarely, but still, 0.42% of segments rated high on TT (ratings of 4 or 5) were also rated high on VP (rating of 4 or 5). 18% of segments were rated as having both some TT (ratings 2-5) and some (ratings 2-5). Similarly, in periods with low vocal activity, a segment could be rated as having both low TT and low VP.

Ratings for TT and VP were dissimilarly distributed across the Likert-scale range, as shown in Table 3. This distributional difference makes direct comparison across segments designated as having varying degrees of TT or VP complicated. In order to compare levels of TT and VP with maximally similar numbers of segments at two levels in both cases, we split TT ratings into “No Turn Taking” (rating of 1) vs. “Any Turn Taking” (ratings 2-5), and VP ratings into “Low Vocal Play” (ratings 1-3) vs “High Vocal Play” (ratings 4-5) levels. Even with this procedure, the TT split yielded a dramatic imbalance, with > 80% of all segments pertaining to the No TT grouping and 82% of the Any TT segments rated 2 (“less than half the time” on TT). On the other hand, VP was very common, with only 8% rated as having no VP, and 55% rated as having high VP, occurring either “more than half the time” or “close to the whole time” (Fig 2). Crosstabulation counts and percent of total segment ratings across TT and VP variables for both Risk groups are provided in Supporting Information (SI), Appendix A, illustrating that the distributions for TT and VP were very similar for both risk groups.

**Table 3.**
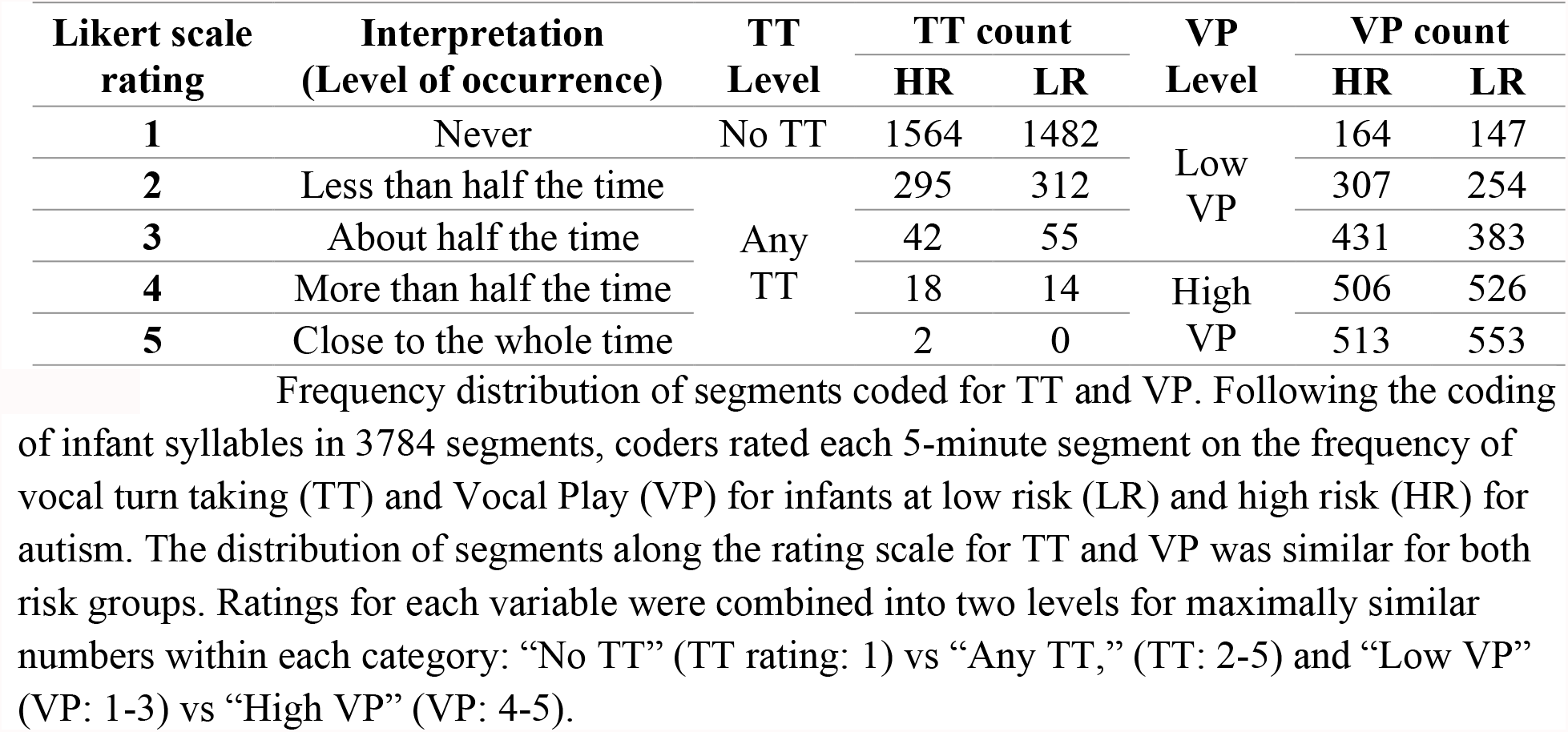
Segments rated for Turn Taking and Vocal Play.

### Coder agreement

Inter-rater agreement was examined for CBRs, TT level, and VP level using a secondary LENA recording dataset coded by 7 of the same graduate student coders following the coding protocol used in the present study and in Oller et al. [27]. The agreement coding for canonical babbling ratios revealed high agreement for both the entire set, with ages ranging across the entire first year (*r* = .89), and for the subset that pertained only to the second half year (*r* = .87), a time period during which CBR varies substantially above 0 across the entire range of ages. Both the questionnaire items yielded far better than chance levels of agreement on the Likert-scale judgments categorized binarily as in the present work (No TT = 1, Any TT = 2-5; Low VP = 1-3, High VP = 4-5) based on Chi square analysis (*p* < .001). For VP there was agreement on 66% of pairings, while for TT there was agreement on 87%, with only fair agreement on kappa (TT = .40, VP = .33). This level of agreement should offer little surprise, given the subjective nature of the judgments. We have been surprised, however, by the power to significantly predict CBR that these blunt measures offer, as will be seen below. See SI, Appendix B for additional methodological rationale and empirical information regarding Coder Agreement.

### Statistical approach

We used Generalized Estimating Equations (GEE) [53] implemented in R for all analyses (see SI, Appendix B for rationale). An expanded version of our results is included in SI Appendix C showing preliminary interaction models across Age, Risk, TT, and VP.

## Results

First, in evaluation of our first hypothesis, we observed mean CBRs across the three ages—7.5 months, 9.5 months, and 12 months—to confirm that our data showed the well-established increase in CBRs during the second half-year of life (Fig 1A). Our results revealed a significant effect of Age at both intervals (7.5 to 9.5 months, *p* < .001, *b* = .04; 9.5 to 12 months, *p* < .001, *b* = .04), evidencing a strong and near linear increase of CBRs over time for data amalgamated across all infants, independent of TT and VP.

**Fig 1.**
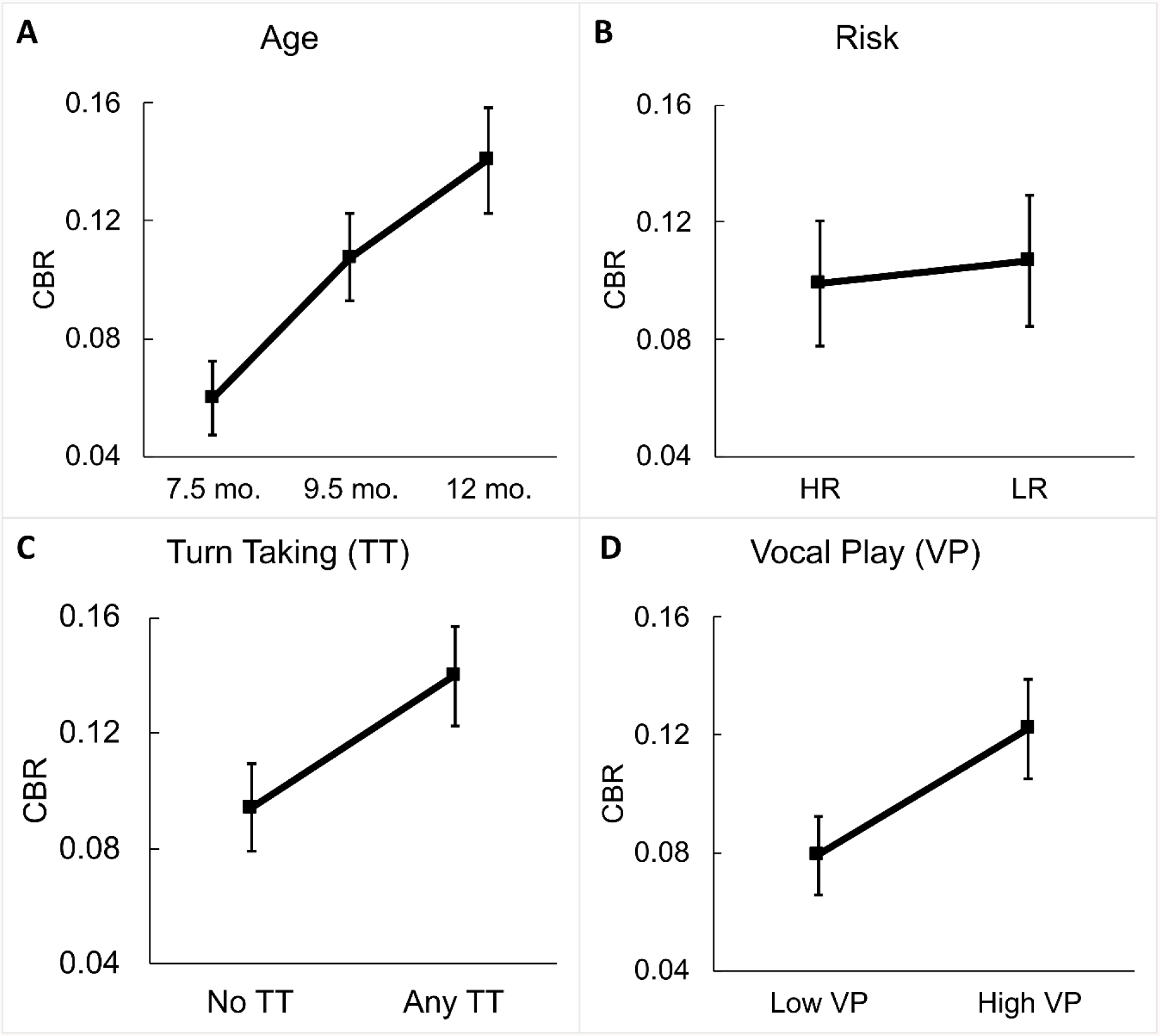
Main effects for Age, Risk, Turn Taking, and Vocal Play. Fig 1A illustrates the significant main effects of Age between 7.5 and 9.5 months (p < .001, b = .04) and 9.5 and 12 months (p < .001, b = .04). 1B shows the non-significant main effect of Risk group (p = .742). 1C presents the significant main effect of Turn Taking, with higher CBRs during segments rated as having any TT compared to those rated as having no TT (p < .001, b = .04). Finally, 1D shows the significant main effect of Vocal Play, with higher CBRs present during segments with high VP compared to segments with low VP (p < .001, b = .06). Standard error (SE) bars are shown.

Considering the second hypothesis, we then compared overall CBRs between infants at low and high risk for autism to explore group-level prelinguistic differences that could potentially indicate later language impairments (Fig 1B). We observed no significant differences in mean CBRs between the two Risk groups (*p* = .742), suggesting no major effect of autism risk status on canonical syllable production in the first year of life.

On hypothesis 3, examining infants’ production of advanced vocal forms across periods with varying levels of turn-taking, we compared infants’ mean CBRs between segments grouped as having No TT (Likert scale rating = 1) and Any TT (Likert scale ratings = 2-5), shown in Fig 1C. We observed a significantly higher mean CBR during Any TT (M = 0.14, SD = 0.17) than No TT (M = 0.09, SD = 0.14) indicating an increased production of canonical babbling during periods with high social vocal activity compared to low social vocal activity (*p* < .001, *b* = .04).

And on hypothesis 4, examining infants’ production of advanced vocal forms across varying levels of vocal play, we compared infants’ mean CBRs between segments grouped as having Low VP (Likert scale ratings = 1-3) and High VP (Likert scale ratings = 4-5), shown in Fig 1D. We again observed a significantly higher mean CBR during High VP (M = 0.12, SD = 0.16) than Low VP (M = 0.08, SD = 0.13) suggesting increased canonical babbling during periods with high endogenous activity compared to low endogenous activity (*p* < .001, *b* = .06).

## Discussion

The present work evaluated canonical babbling ratios (CBRs) in 98 infants (identified as being at low or high risk for autism) across 3784 five-minute segments, selected from all-day recordings in the infants’ homes across the second half-year of life. The recorded segments were coded by a team of highly trained listeners, who determined CBRs and subjective rates of occurrence of vocal turn taking (TT) and vocal play (VP) in each segment. We confirmed that the production of canonical syllables increased across the second half-year of life and found a significantly higher production of canonical babbling during both high TT and high VP. Interestingly, there were no differences observed in CBRs between infants at low and high risk for autism. Our work offers perspective on the role of social and endogenous motivation in the emergence of advanced vocal forms in prelinguistic development, suggesting that both social and endogenous factors are at play.

### Age and autism risk

With respect to our first hypothesis, our findings revealed, of course, the expected significant effect of Age on CBRs, a finding consistent with all prior longitudinal studies of canonical babbling. For our second hypothesis, we predicted that infants with an elevated likelihood for having autistic characteristics (“high risk”, due to the presence of an older sibling diagnosed with ASD) may have reduced social motivation to signal fitness as compared to infants with a typical likelihood for ASD (“low risk,” due to having no siblings or other near relatives diagnosed with autism). However, we observed no main effect of autism risk on CBRs. The finding adds further uncertainty to the already mixed evidence on canonical babbling emergence in both autism diagnosis and risk [48,54,55]. The results support the argument that canonical babbling may be a robust developmental phenomenon across contexts and may be more resistant to cognitive effects of risk than has been expected.

### Social and endogenous factors in protophone production

The present data do, on the other hand, provide novel findings regarding our third and fourth hypotheses. We posited that early language development is driven by both infant social motivation (presumably reflected in infant interest in caregiver vocalizations and in protoconversation) and infant endogenous inclination to produce copious amounts of vocalization, a tendency that appears to have been naturally selected as a signal of fitness. We observed high CBRs across all infants during segments with TT *and* during segments with high VP. The effect of TT was considerable, being equivalent to 2-3 months of growth in CBR, and the effect was even larger for high VP. Thus the results suggest that both social interaction and exploratory vocalization stimulate canonical babbling.

### Social motivation and turn taking

The findings for CBRs during TT (alongside no major differences between risk groups) suggest similar levels of social motivation in both groups, with all infants showing the tendency to produce higher CBRs during segments rated as having any TT compared to those rated as having no TT. The social motivation reasoning behind our predictions was based in the assumption that infants at high risk for autism may present with a greater likelihood for reduced experience of social reward compared to infants at low risk and thus may demonstrate reductions in vocal performance during social interaction [31,32]. These findings suggest robustness of social motivations for infant vocalization. Our hypotheses were based on an expectation of anomalous development in HR infants, assuming social motivation for vocalization may break down in the presence of neurodevelopmental factors negatively affecting social cognition. The results suggest a stronger mechanism where human infant vocal tendencies may have been selected to withstand neurodevelopmental deficits associated with autism. Alternatively, the results might hint at adaptation by parents, who may adjust their interactive styles to be more effective in eliciting canonical babbling from their infants at risk than in the case of parents whose infants are not at risk.

There can be no doubt that humans are highly social. Clearly, early hominins’ relatively large living groups necessitated a high level of social bonding, which created a need for an efficient communication method, resulting in positive selection pressures on the evolution of language [56–58]. Chevallier [31] noted that “social motivation constitutes an evolutionary adaptation geared to enhance the individual’s fitness in collaborative environments” (p. 2). Thus, it is reasonable to assume that precursors to language such as canonical babbling must be robust during development to drive its evolution. Although often reported to be delayed in developmental disorders, including autism [54,59,60], canonical babbling is well-established as a robust stage of development, known to emerge even in infants at risk because of premature birth or very low socio-economic status [61,62]. Our results suggest that the most advanced prelinguistic vocal forms (i.e., canonical syllables) produced during early face-to-face interactions may be robust even in the face of a potentially elevated likelihood for social communication deficits.

#### Measurement vulnerability

One important consideration and potential limitation in our evaluation of social motivations in early infancy relates to the measures we used to assess the sociality of vocalizations. To measure social and exploratory vocal functions, coders were asked to estimate on a Likert scale how often infants engaged in TT and VP for each segment. This subjective measure, obtained immediately after coding for CBR for each segment, can be portrayed as a blunt instrument, subject to only fair inter-observer agreement, but it is founded in the notion that human judgments are the gold standard for any such measure. In addition, this method of obtaining the judgments was convenient and workable within the time constraints of the coding participants. A perhaps more reliable measure would require labeling the social or exploratory function of each utterance individually with repeat-observation (and especially with both audio and video), a measure that requires at least tenfold more time to obtain (see Long et al. (2020) for an analysis using this method). Future studies using this more expensive measure to address the role of TT and VP in infant vocalization are planned. Additional future considerations include examining the role of infant-directed speech (see SI, Appendix E) using methods similar to those employed in this study, as well as taking account of initiation of turn-taking (i.e., whether it is infant-initiated or caregiver-initiated) and Risk to further study potential social motivations of the infant.

TT occurred, according to the coders, in only about 20% of the segments at all (and the vast majority of such segments were rated 2, meaning TT occurred during less than half the segment), a pattern that applied roughly equally to both Risk groups. This low rate of TT surprised us, given that so much of the literature on early language development focuses on protoconversation and its presumable importance in development. The low rate of TT surely imposed a power limitation on the statistical analyses of the effects of TT and its interactions with the other variables in the present work. The fact that the analysis produced robustly significant results even in the face of the power limitation suggests a robust relation between TT and canonical babbling.

### Endogenous activity and independent vocal play

The VP measure was also based on a Likert scale, where coders were asked to judge each segment on how much of the time the infant had engaged in independent, not-socially-directed vocalization (presumably endogenously motivated). Unlike TT, VP was found by the coders to be present in the vast majority of segments, and again this was true of both Risk groups—the plurality of segments having been rated 5 (VP present in close to the whole segment) by the coders for both Risk groups. Our surprise in finding low rates of TT in the all-day recordings is matched by our surprise at the near omnipresence of VP. As in the case of TT, the fact that the analysis produced robustly significant results even in the face of the power limitation imposed by only moderate coder agreement suggests a robust relation between VP and canonical babbling.

Interestingly, a preliminary analysis on VP data revealed a significant difference in CBRs between Risk groups over time, but only during periods of high VP (*p* = .039, *b* = .06, see SI Appendix C for additional analyses).

We presumed that autistic infants may prefer playing with the sensorimotor characteristics of a syllable through repetition (reduplication), while their neurotypical counterparts may be more likely to play with varying aspects pertaining to individual syllables, modifying duration, placement, and various articulatory patterns from utterance to utterance. Thus, producing repetitive physical and acoustic properties of sounds during bouts of VP may be more intrinsically rewarding to infants with autistic characteristics compared to those without them, the latter perhaps tending more to explore phonetic nuances. However, additional research is necessary to confirm these tendencies. Our present work highlights the likely importance of endogenous vocal play as sensorimotor exploration of the vocal mechanism, which may be necessary to learn and master speech production. This is supported by the high frequency of segments identified as having “High VP” and the higher CBRs during segments rated as High VP compared to Low VP.

A potential limitation of this study is that we only evaluated the production of *canonical babbling* as a measure of advanced vocal forms. Future studies should compare variation in infant vocal types based on phonatory characteristics across contexts, infants’ recombination of syllables at the utterance level, and more specific contextual social or environmental information such as response contingency and infant- and adult-register speech characteristics of parents (See SI Appendix D for considerations regarding infant-directed speech).

## Conclusions

The findings observed in the present study offer perspective on potential social and endogenous motivations in the production of advanced vocal forms like canonical babbling in the first year of life. We observed high rates of canonical babbling during high turn taking (as a measure of social vocal activity) *and* high vocal play (as a measure of endogenous vocal activity). This suggests infant babbling is socially interactive only occasionally in the natural environment of the home, while endogenous, exploratory vocalization occurs very often throughout the day. Selection pressure across the evolution of hominins may have played a role in infants’ increased rate of canonical syllables during turn taking to signal fitness to caregivers, and during vocal play, an activity suggesting that exploration of vocalization must have been naturally selected to be pleasurable for human infants. Because endogenous protophones occur so often during the day, it has been speculated that their production must be noticed, even if semi-consciously, by parents as a signal of infant wellness. It seems likely that infant vocal exploration also supports development of the vocal apparatus required to speak. Our results on the risk groups offer support for a potentially robust social motivation in infancy to produce higher rates of canonical syllables during interaction, even in the presence of an elevated likelihood for social communication deficits.

## Acknowledgments

The authors wish to thank the participating families in Atlanta, GA and graduate student coders of this research.

