## Supplemental Material for "Canonical babbling during vocal turn taking and independent vocal play"

### Appendix A

#### TT*VP Crosstabulations by Risk Group

**S1 Table. TT*VP Crosstabulations – LR infants.** Crosstabulation counts and percent of total segments rated for TT (Turn Taking) and VP (Vocal Play) for LR (low risk) infants.

|  | | | VP | | | | | |
| --- | --- | --- | --- | --- | --- | --- | --- | --- |
|  |  |  | **Low VP** | | | **High VP** | | |
|  |  |  | **1** | **2** | **3** | | **4** | **5** |
| TT | **No TT** | **1** | 112 | 205 | 292 | | 406 | 467 |
|  |  | % of Total | 6.0% | 11.0% | 15.7% | | 21.8% | 25.1% |
|  | **Any TT** | **2** | 25 | 37 | 75 | | 101 | 74 |
|  |  | % of Total | 1.3% | 2.0% | 4.0% | | 5.4% | 4.0% |
|  |  | **3** | 8 | 10 | 14 | | 17 | 6 |
|  |  | % of Total | 0.4% | 0.5% | 0.8% | | 0.9% | 0.3% |
|  |  | **4** | 2 | 2 | 2 | | 2 | 6 |
|  |  | % of Total | 0.1% | 0.1% | 0.1% | | 0.1% | 0.3% |
|  |  | **5** | 0 | 0 | 0 | | 0 | 0 |
|  |  | % of Total | 0.0% | 0.0% | 0.0% | | 0.0% | 0.0% |

**S2 Table. TT*VP Crosstabulations – HR infants.** Crosstabulation counts and percent of total segments rated for TT (Turn Taking) and VP (Vocal Play) for HR (high risk) infants.

|  | | | VP | | | | | |
| --- | --- | --- | --- | --- | --- | --- | --- | --- |
|  |  |  | **Low VP** | | | **High VP** | | |
|  |  |  | **1** | **2** | **3** | | **4** | **5** |
| TT | **No TT** | **1** | 144 | 228 | 328 | | 413 | 451 |
|  |  | % of Total | 7.5% | 11.9% | 17.1% | | 21.5% | 23.5% |
|  | **Any TT** | **2** | 13 | 61 | 87 | | 84 | 50 |
|  |  | % of Total | 0.7% | 3.2% | 4.5% | | 4.4% | 2.6% |
|  |  | **3** | 6 | 10 | 13 | | 7 | 6 |
|  |  | % of Total | 0.3% | 0.5% | 0.7% | | 0.4% | 0.3% |
|  |  | **4** | 1 | 8 | 3 | | 1 | 5 |
|  |  | % of Total | 0.1% | 0.4% | 0.2% | | 0.1% | 0.3% |
|  |  | **5** | 0 | 0 | 0 | | 1 | 1 |
|  |  | % of Total | 0.0% | 0.0% | 0.0% | | 0.1% | 0.1% |

#### Definitions of vocal play and turn-taking

The all-day recording method does not at this point include video for a variety of technical reasons that will hopefully be overcome in the not-too-distant future. For the LENA-based work, audio-only (we have fairly good SNR, usually available with LENA), we operate with fairly simple rules of judgment that define VP (also called exploratory vocalization) as follows: for a protophone to be judged as vocal play 1) it must not be deemed to be part of a social interaction; 2) it must not be deemed an attempt to initiate an interaction (as in calling for attention); and 3) it must not be deemed to be an accidental sound (an effort grunt for example). These principles are in accord with the original suggestions about vocal play by researchers from the 70’s and 80’s, in particular Betty Stark [1–3]. By our definitions, TT vocalizations produced by the baby are ruled out as VP, so the judgments of VP and TT are intended to be independent, and indeed the correlation between VP and TT judgments is near 0, and the distributions of segments judged to have high and low TT and VP are extremely different (see Tables S1 and S2 above or Table 3 in the main text).

An infant utterance is judged as consisting of TT if it is perceived as a response to another speaker’s infant-directed speech (IDS) utterance. Note that an utterance simply occurring in the neighborhood of speech of others, without actually being interpreted as responsive to that speech of others, does not qualify as TT according to our definition. In general, listeners treat infant utterances as TT if they begin well after (>200 ms after) the onset of the IDS utterance to which they are interpreted as being responsive, and not longer than 3 seconds after the offset of that IDS utterance. These time values have been assessed empirically [4].

### Appendix B

#### Coder Agreement

**Coding agreement rationale**

Agreement among coders does not, in principle, need to meet a specific criterion of magnitude. Agreement must, however, be significantly better than chance in order for it to provide a possible basis for discerning effects on a parameter such as CBR. If coder agreement is relatively low, the chance of discerning a real difference is more limited. One can think of low agreement as constituting noise in the analysis and as imposing a power limitation. The fact that significant effects of VP (and TT) were found, even though coder agreement was not high (by typical standards) is evidence of real effects, and leaves open the possibly of greater real effects if the observational system can be improved (especially by moment-to-moment coding of infant utterances instead of questionnaire responses at the end of each five min segment) or by more extensive training on the questionnaire items.

In our work, human observation is expected to show variable categorizations across coders for vocal types and social event types, even after careful training. Infant actions are also expected to vary. Both the infant actions and the judgments of them by parents or coders (whom we often portray as simulated parents) must be expected to vary because these factors are surely under selection pressure, which in Darwinian theory require variation, because otherwise there could be no selection on them. In fact, coder agreement levels on these and other of judgments are a source of important data regarding the real distributions of human reactions to events that matter in child development, for example, natural variations in how parents interpret infant actions, whether the infants are fussing or not, whether they are imitating or not, and so on [5,6].

These points run counter to a tradition in psychology that attempts to impose specific criteria for coder agreement levels (or effect sizes or alpha levels) on all research. Such criteria are ultimately unworkable, because they tend to ignore a host of related factors that determine plausible or meaningful agreement levels (or effect sizes or alpha levels). For ethological research (of which the present work is an example), we are obliged to quantify phenomena that are often fuzzy by nature, and yield by nature lower agreement levels among coders than the tradition in psychology often advocates. However well that tradition seems to work in laboratory experimentation (the most common mode of psychological research), it is often impossible to implement such traditions in natural observational research without sacrificing much of the primary data of interest on the grounds that agreement criteria cannot be met.

Still we will seek to impose more extensive training on many of the questionnaire items of our coding approach, including VP and TT, in the future. We also plan to employ, when practicable, coding of each infant utterance in terms of VP and TT in order to cross check the validity of the questionnaire approach, and potentially to provide more powerful measures. At the time we designed the research reported here, we did not anticipate the effects of VP and TT on CBR that were observed. Yet the effects were robust. Had we known that these questionnaire items would reveal the patterns we found, our training emphasis on VP and TT would surely have been higher.

The coding we do is time-consuming and costly, so there are limits to what can be added, even now that there is more reason to focus on the VP and TT parameters. The questionnaire items were instituted because they allowed low-cost initial assessments of parameters that so far cannot be assessed by automated tools. They would even impose considerable costs (beyond our budgets) on *real-time human coding* of individual infant utterances on TT and VP.

**Coder agreement data for the present study**

Here we review three types of coder agreement data relevant to the present study. We also refer the reader to prior papers (especially their Supplementary Materials sections) in order to obtain a fuller view about our coding training and coder agreement procedures [7–9].

1. The data presented on coder agreement in the main text pertain to the following particular study, which was conducted *after* all the coding was done that was used to compute the results reported here and in the main text. Inter-rater agreement was examined for CBRs, TT level, and VP level using a secondary LENA recording dataset coded by 7 of the same graduate student coders following essentially the same coding protocol used in this study. The 5-minute segments that had already been coded—each by one of the 7 individuals—came from a set of over 1000 such segments randomly selected from the all-day recordings of eight infants at each of six ages across the first year of life. >380 of these segments had been coded in the very same way as in the present study, with determination of CBR, TT, and VP. A subset of 212 of these segments was semi-randomly selected to be assigned for a second pass of agreement coding, where the agreement coder would always be a different individual from the one who had provided the original coding. The number 212 was based on available coder time and the desire for a large enough sample to yield trustworthy agreement data.

Every one of the 7 agreement coders was assigned to at least 5 segments that had originally been coded by each of the other 6 coders. In addition, all agreement coders were assigned to at least 5 segments from each of the 8 infants. Finally, all the ages of infants were included for assignments to each of the agreement coders for at least 5 segments. The agreement coding was conducted blind, in the sense that no coder knew who had originally coded the segments assigned to them in the agreement phase, nor were they supplied with information about age or identity of the infants.

We resolved to report on this agreement study in the main text on the grounds that this was the mostly tightly designed and carefully implemented of all our agreement studies in recent years, using the methods of training and implementation as in the present work.

2. A second study of agreement was conducted after the coding had been completed for nearly half the data reported in the Results of the present work. The agreement study involved 10 of the 16 coders on a subset of the very sample of recordings that is reported on in the Results of the present work. Each of 9 agreement coders was assigned to recode 5-min segments (there were 346 selected for the agreement study) that had been completed months before by the 10 coders. The assignments were made at random and in >95% of cases, the coders were assigned to segments that had been originally coded by a different individual. The number of agreement assignments given to each coder varied (mean = 38.3 segments, sd = 17.7) because the assignments were based on how many hours of coding were available for each of them given their assignment schedules, which were individualized to account for coder classroom and clinic requirements and to the sizes of their original recording assignments, which also varied due to differences in the number of recordings available for the infants to which each of them had been assigned.

Agreement among the coders was comparable to the outcomes for the 7-coder agreement study reported in the main text. For correlations of the original and agreement coding: regarding noncanonical syllables, canonical syllables, CBR, VP and TT respectively, *r* = .65, .86, .82, .53 and .40. All the correlations were significant at *p* < .0001. As in the case of the 7-coder study, the VP and TT measures yielded lower agreement than the other parameters, but all the *r*’s were far above chance level. Reclassifying the 1-5 scales so that they matched the High vs Low VP and No vs Any TT categorizations used in the statistical analyses, 70% of VP judgments were in agreement between the original and agreement coders and 67% of the TT judgments were in agreement. Chi-square values for 2x2 tables of VP and TT judgments also showed that agreement was significantly greater than chance in both cases (*p* < .0001).

3. Another assessment of agreement among the coders was made to include the 6 coders out of the total 16 who were not involved in either of the specific agreement studies. We will call these the NAg (non-agreement study) coders to contrast them with the Ag coders (the ones who had been in the specific agreement studies). This agreement assessment is based on the data we acquired in the final period of training, which was conducted across intensive sessions of presentations of examples and practice coding using the laboratory coding software over a six-week period.

In these cases the coders could all be referenced to each other on independent coding of the same 5-min segments from the set we call “bundle 0”; that set was provided by the Marcus Autism Center author, Ramsay, from among recordings of infants recruited to the same project, but that had not been specifically selected for the study. From bundle 0 we have developed a training set for which we have considerable data on >40 coders over the past 6 years. Using that set we can assess agreement levels on the parameters assessed in the present study for the 6 NAg coders with respect to 6 Ag coders. Since the agreement levels we now report do not result from a designed agreement study, we have adapted the comparisons so that all 6 NAg coders are paired for comparison with all 6 Ag coders, yielding 36 pairings. We take the average correlation across all these pairings for each parameter. Regarding noncanonical syllables, canonical syllables, CBR, VP and TT respectively the average *r* = .76, .88, .82, .34 and .59.

Appendix B

#### Statistical Analysis

We used Generalized Estimating Equations (GEE) implemented in R for all analyses. First we analyzed main effects and interactions of Risk, Age, and TT on infant CBRs, then Age Risk, and VP on infant CBRs. We also tested a simplified model including only Age and Risk. Finally we tested independently for main effects of all four independent variables.

GEE analyses are an advanced form of modeling providing a non-parametric alternative to generalized linear mixed models for estimating within-subject covariance and population-averaged model parameters [10]. GEE has advantages over other mixed models frameworks especially in cases where data across conditions and from participants are intercorrelated and where numbers of observations per participant or condition vary. Another advantage is that the GEE approach requires no normality assumption. A GEE analysis is appropriate here because this is a longitudinal dataset with an unequal number of observations on infants, number of recordings per Age and Risk group, and number of observations of TT and VP ratings within each level.

### Appendix C

#### Expanded Results

We ran three GEE models evaluating interactions and main effects for 1) TT, Age, and Risk, 2) VP, Age, and Risk, 3) Age and Risk, and we ran a fourth GEE model on main effects only for 4) Age, Risk, TT, and VP.

**Turn Taking, Age, and Risk**

1. Based on predictions derived from the social motivation theory, we predicted higher CBRs in low-risk (LR) infants during segments with high turn taking but no such pattern in high-risk (HR) infants. However, the results (Table S3) did not confirm the hypothesis (*p* = .144). The mean CBR for HR and LR infants was quite similar for segments with no TT but showed a complex relation between Risk and Age (though not significantly) for segments with any amount of TT.

**S3 Table. Turn Taking, Age, and Risk interaction model.** Full interaction GEE model for CBR with Age group (7.5 to 9.5, and 9.5 to 12 mo.), Risk Group (HR vs LR), and Turn Taking as a factor (No TT vs Any TT).

| Variable | Effect size (*b*) | SE | *p* |
| --- | --- | --- | --- |
| TT (No vs Any) | 0.02 | 0.02 | 0.347 |
| Risk (LR vs HR) | -0.02 | 0.01 | 0.111 |
| Age (7.5 to 9.5 mo.) | 0.06 | 0.01 | < .001** |
| Age (9.5 to 12 mo.) | 0.02 | 0.02 | 0.121 |
| TT * Risk | 0.04 | 0.03 | 0.144 |
| TT * Age (7.5 to 9.5 mo.) | 0.01 | 0.02 | 0.812 |
| TT * Age (9.5 to 12 mo.) | 0.02 | 0.02 | 0.426 |
| Risk * Age (7.5 to 9.5 mo.) | 0.05 | 0.02 | 0.004** |
| Risk * Age (9.5 to 12 mo.) | 0.03 | 0.02 | 0.075 |
| TT * Risk * Age (7.5 to 9.5 mo.) | -0.04 | 0.04 | 0.368 |
| TT * Risk * Age (9.5 to 12 mo.) | -0.05 | 0.03 | 0.136 |

In the full GEE model, we found no main effect of TT, that is, no significant difference in CBRs between segments rated as having No vs Any TT (*p* = .347). Differences in CBRs between Risk groups were also non-significant (*p* = .111), with somewhat higher CBRs in the LR group. In the same model, the main effect for CBR from 7.5 to 9.5 months of age was highly significant (*p* < .001, *b* = .06), but differences from 9.5 to 12 months were not (*p* = .121), reflecting the fact that CBRs went up more from 7.5 to 9.5 months than they did from 9.5 to 12 months.

The results did not show significant two-way interactions between TT and either of the Age group comparisons: 7.5 to 9.5 months (*p* = .812) or 9.5 to 12 months (*p* = .426). There was, however, a significant interaction between Risk and Age for 7.5 to 9.5 months (*p* = .004, *b* = .05); CBRs increased in HR infants to a greater extent between 7.5 and 9.5 months then in LR infants across these two ages. This difference was reversed from 9.5 to 12 months such that LR infants (*p* = .075) showed a greater increase than HR infants in that age interval, an interaction that approached statistical significance. No significant three-way interactions were observed in this model. Fig S1 provides graphic illustration of the results presented in the full model for Age, Risk, and TT level.

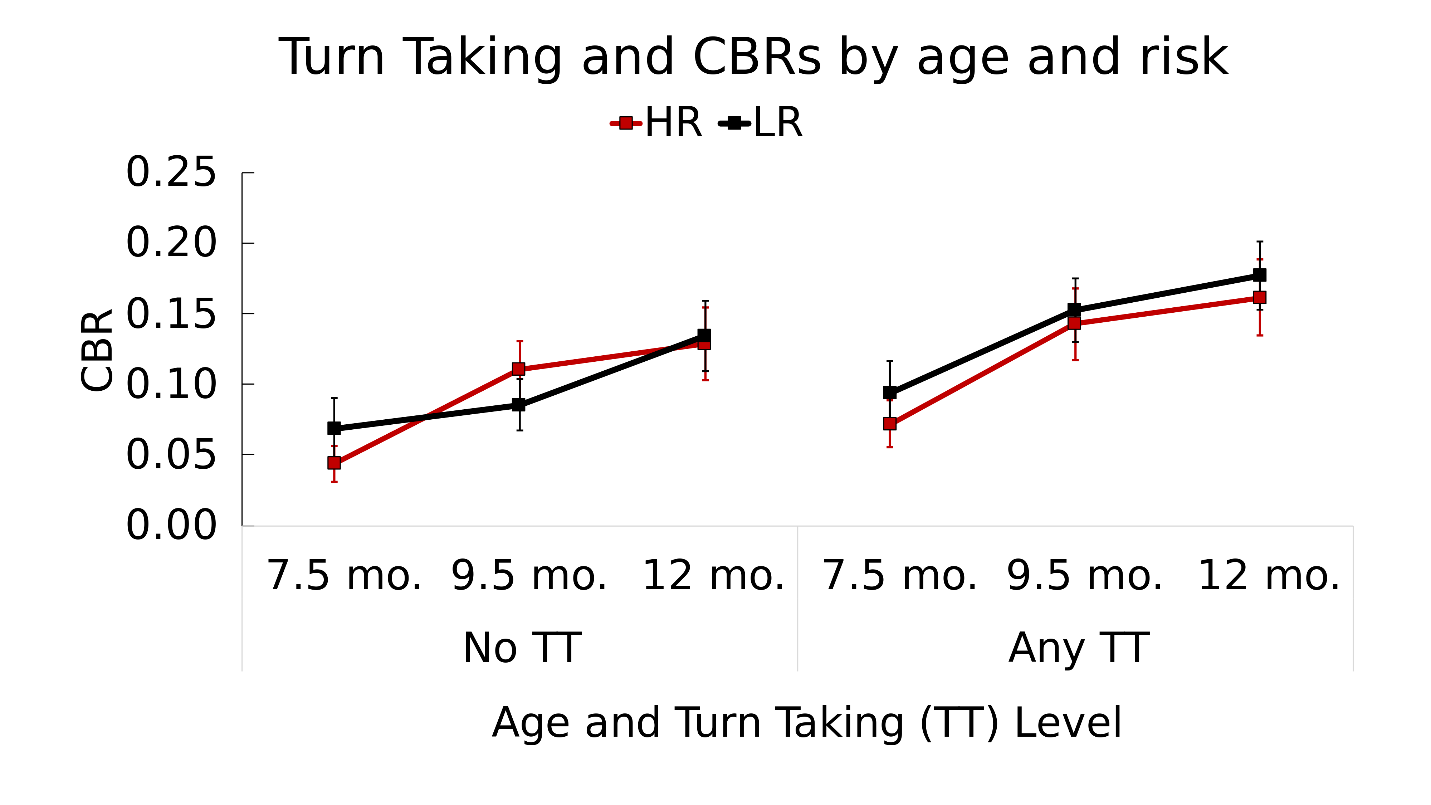

**S1 Fig. Canonical babbling by Age, Risk, and Turn Taking level.** Canonical babbling ratios of infants at high-risk (HR) and low-risk (LR) for autism during segments with no vs any turn taking (TT) across three age ranges, 6.5-8.49 (7.5 months), 8.5-10.49 (9.5 months), and 10.5-13 (12 months) months. CBR was significantly higher from 7.5 to 9.5 months (p < .001, b = .06), and there was a significant two-way interaction of Risk and Age again between 7.5 and 9.5 months (p = .004, b = .05). There were no significant interactions including TT as a variable, including the three-way interactions of Age, Risk, and TT level. The values presented in the figure were computed from the raw data with means and SEs weighted for the number of infants who contributed data in each Risk group at each Age.

##### Vocal Play, Age, and Risk

In the full GEE model for Age, Risk, and VP (Table S4) we found several significant effects not observed in the model for Age, Risk, and TT.

**S4 Table. Vocal Play, Age, and Risk interaction model**. Full interaction GEE model for CBR with Age (7.5, 9.5, and 12 mo.), Risk (HR vs LR), and Vocal Play (Low VP vs High VP).

| Variable | Effect size (*b*) | SE | *p* |
| --- | --- | --- | --- |
| VP (Low vs High) | 0.09 | 0.01 | < .001** |
| Risk (LR vs HR) | 0.00 | 0.02 | 0.800 |
| Age (7.5 to 9.5 mo.) | 0.03 | 0.01 | 0.023* |
| Age (9.5 to 12 mo.) | 0.05 | 0.02 | 0.001** |
| VP * Risk | -0.03 | 0.02 | 0.021* |
| VP * Age (7.5 to 9.5 mo.) | -0.06 | 0.02 | 0.001** |
| VP * Age (9.5 to 12 mo.) | -0.04 | 0.02 | 0.059 |
| Risk * Age (7.5 to 9.5 mo.) | 0.02 | 0.02 | 0.426 |
| Risk * Age (9.5 to 12 mo.) | -0.01 | 0.02 | 0.679 |
| VP * Risk * Age (7.5 to 9.5 mo.) | 0.06 | 0.03 | 0.063 |
| VP * Risk * Age (9.5 to 12 mo.) | 0.06 | 0.03 | 0.039* |

*2. Predicted interaction of Risk and VP*: Based on predictions derived from the social motivation theory, we predicted *an increase in CBRs in HR infants from segments with low to high VP, and a lesser increase or no increase from low to high VP for LR infants*. There was indeed a significant interaction between VP level and Risk group (*p* = .021, *b* = -.03), but the direction of the effect was the opposite of that predicted. CBRs of LR infants increased to a greater extent from low to high VP than CBRs of HR infants. Based on calculations for Fig S2, CBRs at low VP were comparable (HR = .079, LR = .080), while those at high VP differed more, favoring the LR group (HR = .119, LR = .124).

There was a highly significant main effect of VP, corresponding to a higher overall mean CBR produced by all infants during high VP compared to low VP (*p <* .001, *b* = .09). As with the full TT model, we observed no significant difference between Risk groups in the full VP model. There was, however, a significant effect of Age at both levels in the full VP model, with CBRs significantly increasing between ages 7.5 and 9.5 months, (*p* = .023, *b* = .03) and between ages 9.5 and 12 months, (*p* = .001, *b* = .05).

There was a significant two-way interaction for CBR between VP level and Age for 7.5 to 9.5 months (*p* < .001, *b* = -.06); this interaction reflects the fact that CBRs differed more between high VP and low VP at 9.5 than at 7.5 months. The difference between VP level and Age for 9.5 to 12 months approached significance (*p* = .059), and the effect was in the opposite direction, namely CBRs differed less for high VP vs low VP at 12 than at 9.5 months. There were no differences between Risk and Age at either age comparison.

There was a significant three-way interaction between VP level, Risk, and Age for ages 9.5 and 12 months (*p* = .039, *b* = .06). The three-way interaction for Risk, VP level, and Age at 7.5 and 9.5 months approached significance (*p* = .063). Fig S2 provides a graphic display of the effects found with the second model and helps illustrate the nature of the three-way interactions. The data from segments rated as having high VP (right-hand panel) suggest a tendency of CBR to grow rapidly from 7.5 to 9.5 months in the HR infants, but to grow much less rapidly in the LR infants. The opposite growth pattern (LR more rapid, HR less rapid) is seen from 9.5 to 12 months. No such differentiation is observable in the left panel. Thus, the data suggest the LR and HR infants show very different patterns of growth in CBR with age, but only in cases of high VP.

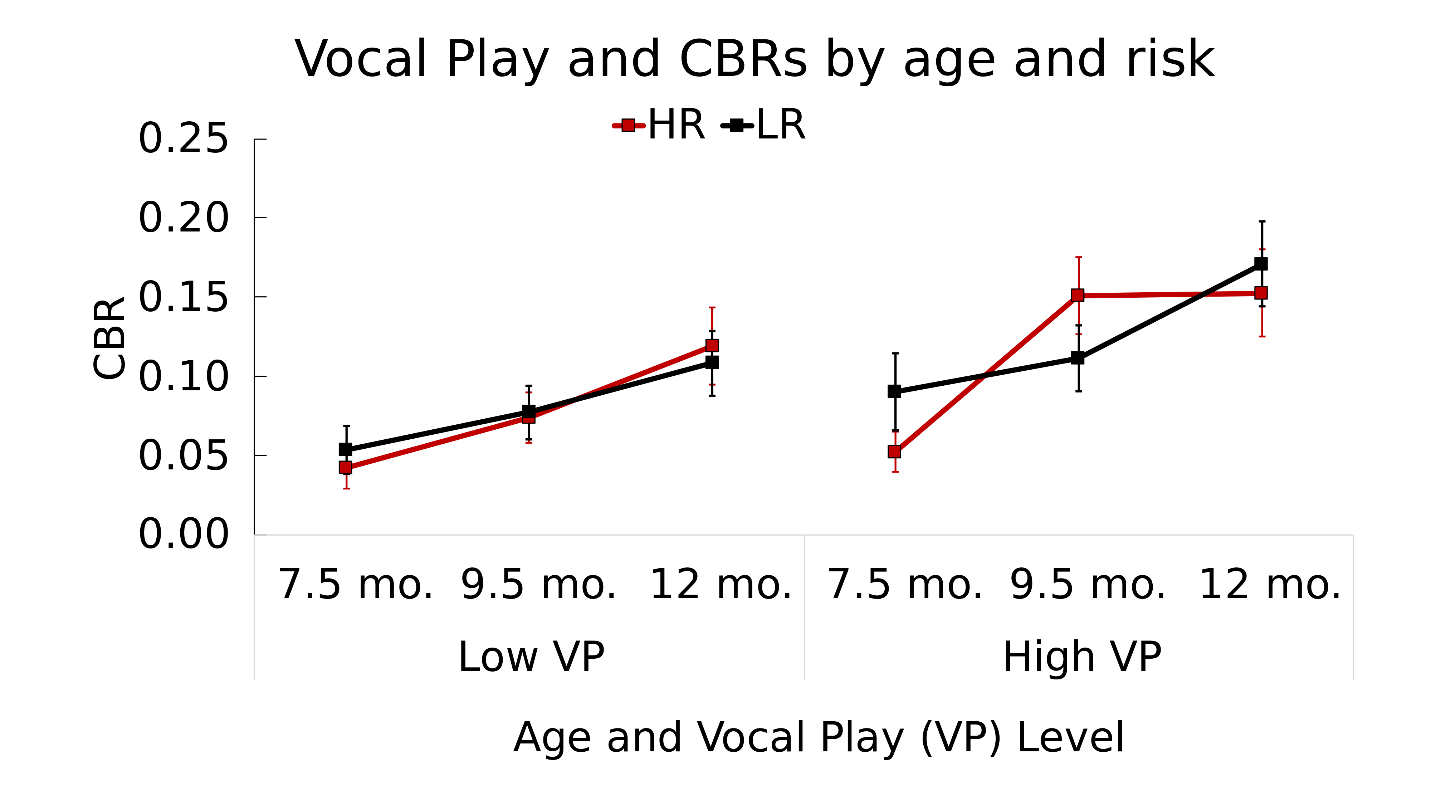

**S2 Fig. Canonical babbling by Age, Risk, and Vocal Play.** Canonical babbling ratios (CBRs) of infants at high risk (HR) and low risk (LR) for autism in segments with low vs high vocal play (VP) across three age ranges, 6.5-8.49 (7.5 months), 8.5-10.49 (9.5 months), and 10.5-13 (12 months) months. In this model, there was a significant effect of Age for both 7.5 to 9.5 months (p = .023, b = .03) and 9.5 to 12 months (p = .001, b = .05). A significant interaction occurred between Risk and VP level (p = .021, b = -.03) and Age and VP level at 7.5 to 9.5 months (p < .001, b = -.06), with the interaction approaching significance for ages 9.5 to 12 months (p = .059). The three-way interaction among VP level, Age, and Risk was significant for ages 9.5 to 12 months (p = .039, b = .06), and approached significance for 7.5 to 9.5 months (p = .063). As in the case of Fig S1, the values presented here were computed from the raw data with means and SEs weighted for the number of infants who contributed data in each Risk group at each Age. Standard error (SE) bars are shown.

##### Age and Risk

3. Based on the preponderance of prior research in autism, we predicted that *CBRs of* *LR infants would increase to a greater extent across the three ages than CBRs of HR infants*. The results did not conform simply to the prediction. In fact CBRs for HR infants rose *more* in the first age interval (from 7.5 to 9.5 months, ~.067 CBR units) than for LR infants (~.015), while they rose *less* in the second interval for HR infants (~.010) than for LR infants (~.065). These patterns corresponded to a significant interaction of Risk by Age at the first interval (7.5 to 9.5 months, *p* = .017, *b* = .04), but a non-significant interaction of Risk by Age at the second interval (9.5 to 12 months, *p* = .192).

Table S5 presents the full GEE model comparing Age and Risk groups. There was a significant main effect for both Age intervals (7.5 to 9.5 months, *p* < .001, *b* = .06; 9.5 to 12 months, *p* = .047, *b* = .03), suggesting an overall increase in CBRs over time, as expected. As in the prior models, there was no significant difference between Risk groups (*p* = .319).

**S5 Table. Age and Risk interaction model.** GEE interaction model for CBR with Age (7.5, 9.5, and 12 mo.) and Risk (HR and LR) only.

| Variable | Effect size | SE | *p* |
| --- | --- | --- | --- |
| Age 7.5 to 9.5 mo. | 0.06 | 0.01 | < .001** |
| Age 9.5 to 12 mo. | 0.03 | 0.01 | 0.047* |
| Risk | -0.02 | 0.02 | 0.319 |
| Risk * Age (7.5 – 9.5 mo.) | 0.04 | 0.02 | 0.017* |
| Risk * Age (9.5 – 12 mo.) | 0.02 | 0.02 | 0.192 |

Fig S3 illustrates these data, showing CBRs of LR infants increased only slightly in the first age interval and a much larger increase in the second interval. Conversely, CBRs in the HR group increased much more in the first interval than in the second. Comparing this interaction with the data in Figs S1 and S2 offer perspective. In Fig S1 (TT model), Risk and Age interacted such that the greater growth of CBR for HR infants in the first age interval applied primarily to the circumstance of No TT, although the three-way interactions corresponding to this observation were not significant. In Fig S2 (VP model), Risk and Age interacted such that the greater growth of CBR for HR infants in the first age interval applied primarily to the circumstance of high VP, and the three-way interactions corresponding to this observation were significant for the first interval and approached significance for the second.

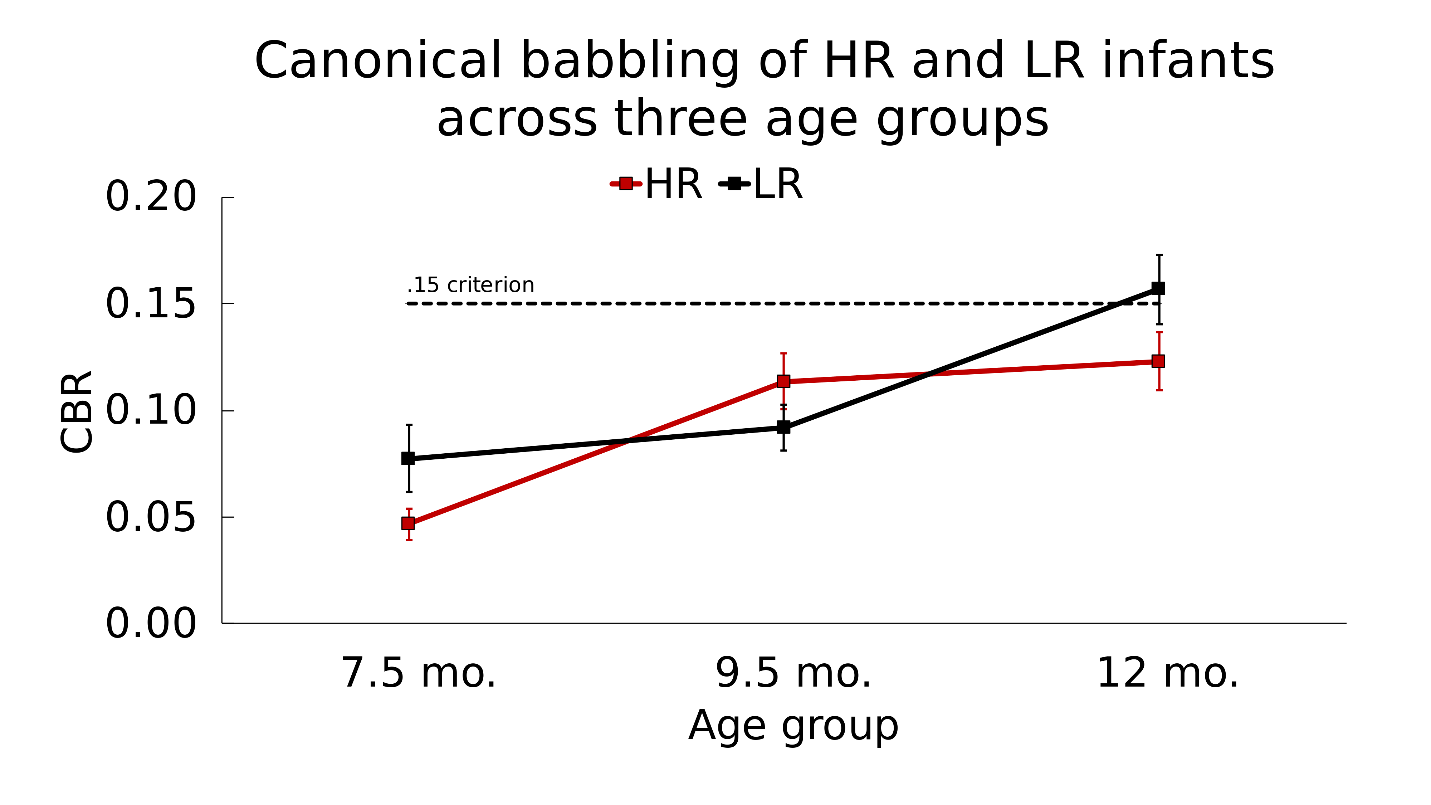

**S3 Fig. Canonical babbling ratios by Age and Risk.** Canonical babbling ratios of infants at high-risk (HR) and low-risk (LR) for autism across three age ranges, 6.5-8.49 (7.5 mo.), 8.5-10.49 (9.5 mo.), and 10.5-13 (12 mo.). Overall, we found a significant interaction of Risk by Age for the first interval (7.5 to 9.5 months, p = .017), with CBRs rising much faster for HR infants than LR infants. The pattern was reversed, but not significantly in the second interval. Standard error (SE) bars shown.

##### Main Effects

In a GEE model analyzing main effects only (Table S6), we found a significant effect of Age at both intervals (7.5 to 9.5 months, *p* < .001, *b* = .04; 9.5 to 12 months, *p* < .001, *b* = .04), evidencing a strong and near linear increase of CBRs over time for data amalgamated across the Risk groups and independent of TT and VP. There was also a significant effect for both TT (*p* < .001, *b* = .04) and VP (*p* < .001, *b* = .06). The effect sizes, reflected in the *b* values from the GEE analysis, can be placed in perspective by considering that TT had an effect roughly of the same magnitude as 2-3 months of growth in CBR, and that VP had an even larger effect.

**S6 Table. Main effects for Age, Risk, TT, and VP.** Main effects model for Age (7.5, 9.5, and 12 months), Risk (LR and HR), Turn Taking (TT) level (No TT vs Any TT), and Vocal Play (VP) level (Low VP vs High VP).

| Variable | Effect size | SE | *p* |
| --- | --- | --- | --- |
| Age 7.5 to 9.5 mo. | 0.04 | 0.01 | < .001** |
| Age 9.5 to 12 mo. | 0.04 | 0.01 | < .001** |
| Risk | 0.004 | 0.01 | 0.742 |
| TT | 0.04 | 0.01 | < .001** |
| VP | 0.06 | 0.01 | < .001** |

The magnitude of the significant effects by Cohen’s *d*, computed from the raw data—with means and SEs weighted for the number of infants who contributed data in each Risk group at each Age—was 0.29 (small) for both TT and VP. The Age effect size was 0.36 (small) for the first interval, 0.21 (small) for the second, and 0.55 (medium) for a comparison of 7.5 months with 12 months. There was no main effect of Risk (*p* = .742). Fig S4 displays these main effects, including significantly higher CBRs during both any TT and high VP compared to periods of no TT and low VP, respectively.

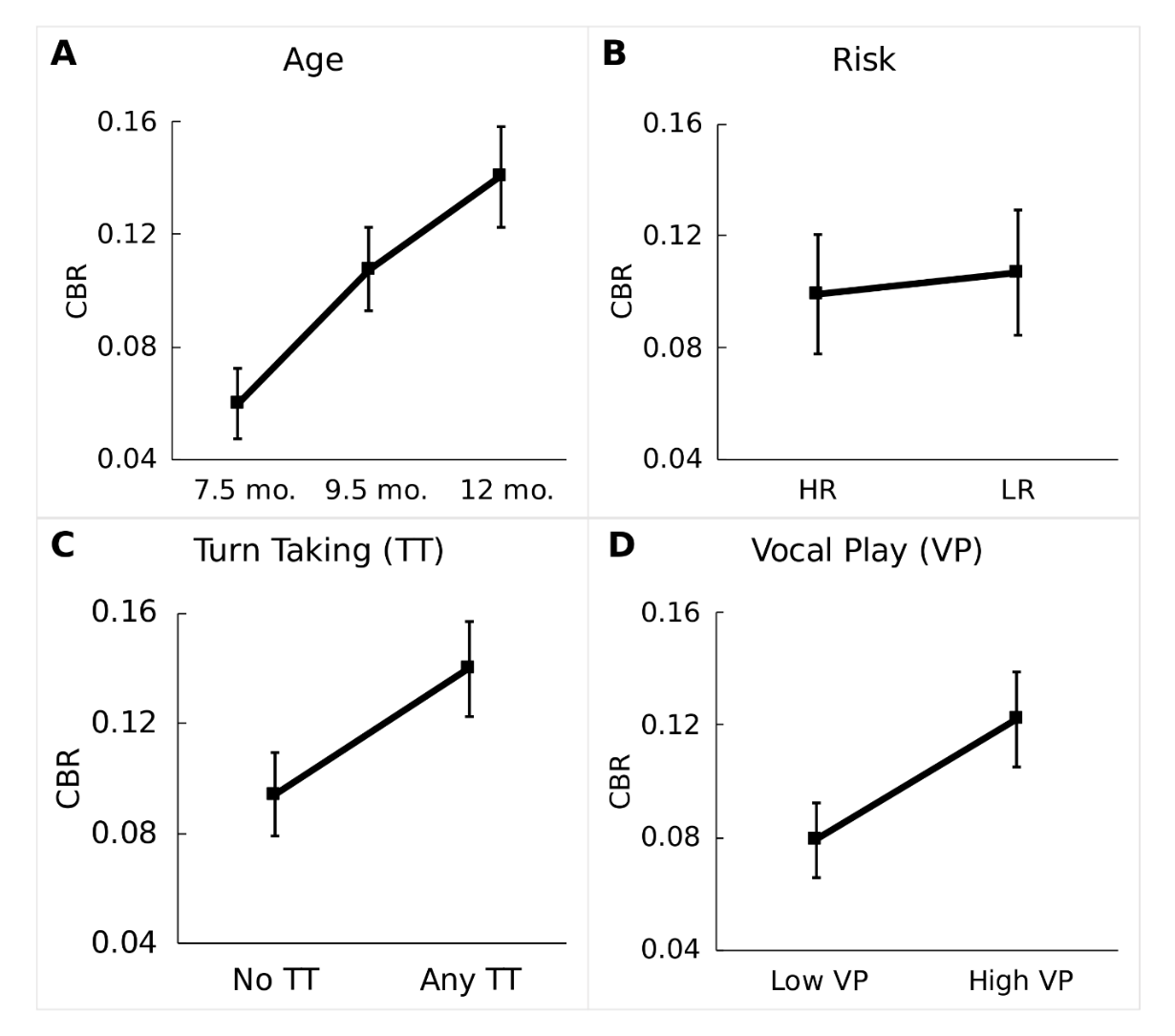

**S4 Fig. Main effects for Age, Risk, Turn Taking, and Vocal Play.** Fig S4A illustrates the significant main effects of Age between 7.5 and 9.5 months (p < .001, b = .04) and 9.5 and 12 months (p < .001, b = .04). S4B shows the non-significant main effect of Risk group (p = .742). S4C presents the significant main effect of Turn Taking, with higher CBRs during segments rated as having any TT compared to those rated as having no TT (p < .001, b = .04). Finally, S4D shows the significant main effect of Vocal Play, with higher CBRs present during segments with high VP compared to segments with low VP (p < .001, b = .06). Standard error (SE) bars are shown.

### Appendix D

#### Considerations Regarding Infant-Directed Speech

The literature on early language suggests infant-directed speech (IDS) may also influence the emergence of canonical babbling, as previous research has highlighted the effects of social interaction on babbling [11–13] and conversely, the effects of babbling on caregiver speech during interaction [14]. During our analyses, we ran a secondary main effects model including IDS as a variable. Our coding protocol also included counts of both infant- and other-directed speech (i.e., speech between two adults) in each segment, affording the opportunity to compare counts or proportions of IDS to CBRs. We found a significant effect of IDS on CBRs (*p* = .034, *b* = -.0004), but notably, this effect was extremely small and negative. Furthermore, the correlation between canonical babbling ratios and total IDS showed a weak, negative correlation (*r* = -.02). In spite of these seemingly negative initial findings, we continue to believe IDS is a variable worth exploring further as an influence on canonical babbling. We will have data on IDS both as a continuous variable based on the coded amount of IDS and as a categorical factor (i.e., Low vs High IDS or No vs Any IDS) based on questionnaire judgments of the same type as explored for TT and VP in the present research. Based on these data we plan to examine IDS effects on canonical babbling more explicitly in a future paper.
